# Genome-wide transcriptome and translatome analyses reveal the role of protein extension and domestication in liver cancer oncogenesis

**DOI:** 10.1101/2021.02.10.430565

**Authors:** Nima Wang, Dalei Wang

**Affiliations:** The First Affiliated Hospital, College of Medicine, Zhejiang University, Hangzhou, China

**Keywords:** liver cancer, translatome, protein extension, domestication, natural selection

## Abstract

One gene could be transcribed to different RNA isoforms, and then produce various forms of protein sequences. This mechanism largely diversifies the cellular pool and allows natural selection to select from a wider range of substrates. Most of the deleterious changes should be either purged or only be observed in patients with deficiencies or diseases. In the cancer field, the “intra-gene” changes between tumor and normal tissues such as the alternative splicing, stop codon read-through, or protein domestication could not be captured by differential expression analyses. In this work, we collected public transcriptome and translatome data from ten patients with liver cancer, and performed genome-wide comparison on the stop codon read-through and protein domestication events. Both events could diversify the proteome without changing the genome sequence. Surprisingly, we found that compared to normal tissues, the tumor tissues globally have significantly higher occurrence of stop codon read-through events. Similarly, the translation signals of non-coding repetitive elements (protein domestication) are elevated in tumor samples. These read-through and domestication events show limited overlapping across the ten patients, suggesting the randomness of the occurrence. It also indicates that these tumor-specific read-through and domestication events should be deleterious, and should be purged by natural selection if they are not collected timely. Our work manifests the role of protein extension and domestication in liver cancer oncogenesis, and adds new aspects to the cancer field.

## INTRODUCTION

With the single-cell sequencing technique becomes more and more popular in the cancer field (Bagger and Probst, 2020), the differential expression analyses between tumor and normal samples seem to be the golden standard of finding the key genes participating in the oncogenesis (Liu et al., 2020; Tieng et al., 2020). However, when talking about one “gene”, we do not refer to only one sequence. One gene could have different transcription start/termination sites (Soong et al., 1994), translation start/termination sites (Kochetov et al., 2005), alternative splicing patterns (Liu et al., 2015), nucleotide modifications (Chandola et al., 2015), protein modifications (Fung and Liu, 2018), and even the switch between coding and non-coding genes, termed domestication (Murray and Murray, 2016). All these factors and their combinations result in the endless isoforms of transcripts and protein sequences, even they are produced from the same gene. Thus, one gene does not only represent one sequence. Traditionally, the differential expression analysis between tumor and normal samples is over-simplified, and it may only find out the genes with very strong functional impact. For the majority of genes that play minor roles in the oncogenesis, more delicate comparisons are needed to determine whether the splicing patterns (or any other mechanisms that change the final protein isoforms) are different between tumor and normal samples. There is compelling incentive for researchers to dig deeper into the molecular changes in oncogenesis. The general scope of this work is to stress the necessity of more detailed insight into the changes at molecular level when studying cancer.

Traditional cancer studies begin with finding key genes (like oncogenes and tumor suppressor genes) responsible for tumor development and oncogenesis. However, instead of particular genes, the oncogenesis could also be triggered by particular mechanisms. For example, any disruptions to the cellular homeostasis could potentially contribute to oncogenesis. The global up- or down-regulation of biological processes (DNA replication, proofreading, repair, mRNA transcription, translation, TE transposition), unbalanced cellular components, and abnormal molecular functions would definitely destroy the steady state of cells (Magistri et al., 2015; Okada and Brennicke, 2006; Sbodio et al., 2016). In the comparison between tumor and normal samples, apart from checking gene by gene, it is equally necessary to check mechanism by mechanism.

In this study, we focus on two mechanisms termed stop codon read-through (protein extension) and protein domestication (from non-coding to coding) (Figure 1). Both process diversify the protein pool. The stop codon read-through is a random process where ribosomes do not stop at the usual stop codons and go on to translate the 3-prime untranslated regions (3′UTR). Normal genes should have ribosome footprints in coding regions (Figure 1A) while the read-through genes have ribosome footprints detected in 3′UTRs (Figure 1B). The extended ribosome signals will be translated into extended proteins. Although read-through is random in most cases, it may also be affected by sequential features and mutations on stop codons. Changes in global read-through probabilities would affect the entire ribosome density in 3′UTRs. The protein domestication process occurs when the non-coding transposable elements (TE) somehow acquire coding ability. Normal TEs just jump in the genome and have nothing to do with the translation machines (Figure 1C) while the domesticated TEs have bona fide translation signals detected in the coding regions (Figure 1D). The global change in protein domestication would also be reflected by the amount of ribosomes translating the noncoding TE.

**Figure 1.**
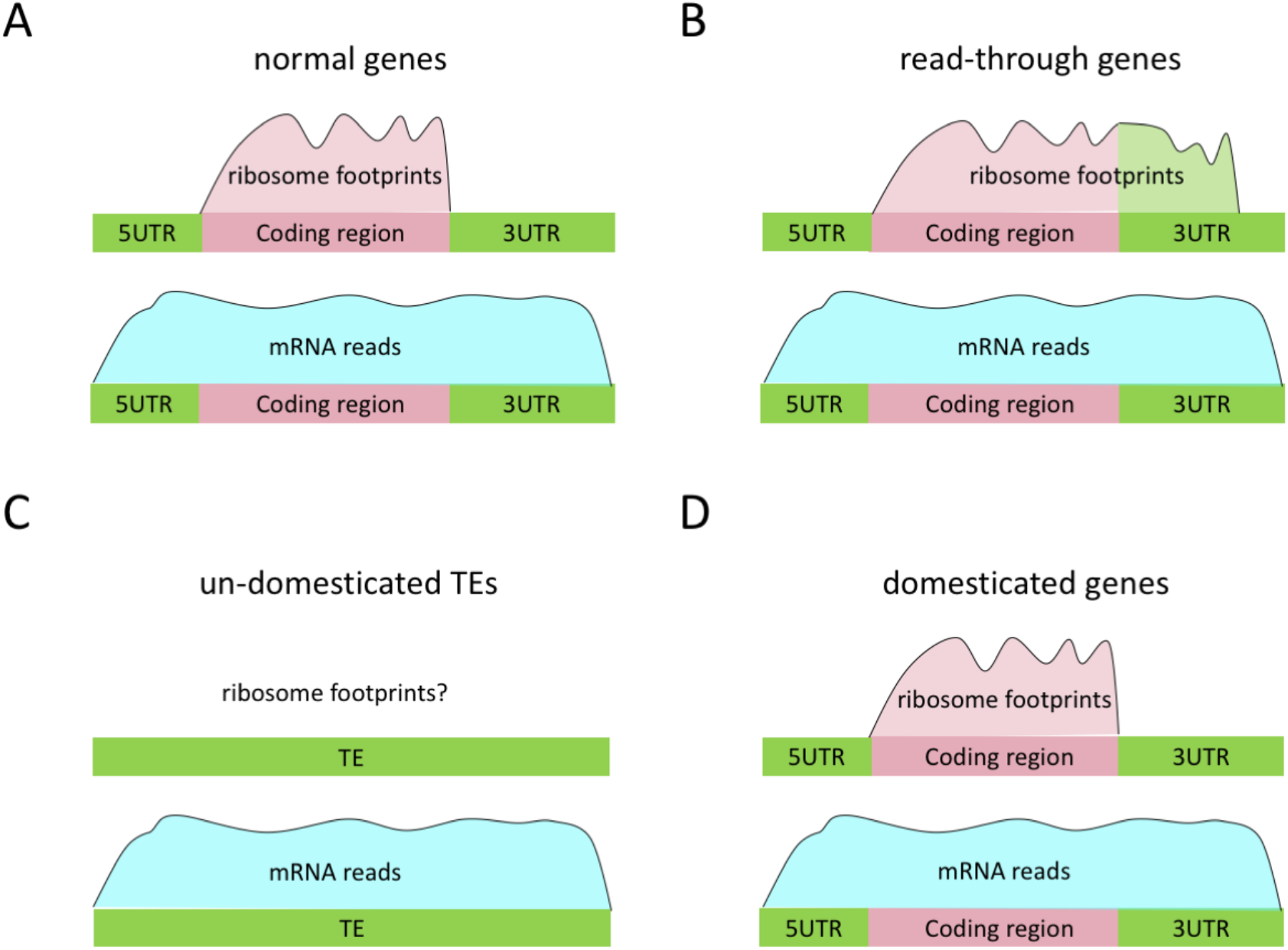
Diagrams telling the readers what stop codon read-through (protein extension) and TE domestication means. (A) For normal genes, only the coding region is covered by ribosome footprints, but the whole gene is covered by mRNA reads. (B) For stop codon read-through genes, both coding region and 3′UTR are covered by ribosome footprints. (C) For un-domesticated TEs, they should not be translated and have no ribosome footprints. (D) For domesticated TEs, the coding region can be translated and covered by ribosome footprints.

We aim to test whether the extent of protein extension and domestication is different between tumor and normal samples. From an understandable aspect of biology, the domestication events represent “from zero to one” while the protein extension events represent “from one to one point five or more”. To accomplish our aim, we collect the transcriptome and translatome data from ten patients suffering from liver cancer (Zou et al., 2019). If these two mechanisms are irrelevant to liver cancer oncogenesis, then we expect to observe similar degrees of protein extension and domestication events. If these mechanisms really contribute to oncogenesis, then we should observe different extent of read-through and domestication events. What’s more, if these tumor-specific changes are deleterious and should not exist in normal tissues, then we should see a skewed frequency spectrum among the ten patients, where many of these tumor-specific read-through or domestication events are singletons, that is, nearly eliminated.

We are going to test our hypothesis among the ten patients suffering from liver cancer. Our results generally support the notion that the and protein extension and domestication mechanisms are correlated with liver cancer oncogenesis. Stop codon read-through ratios and the occurrence of protein domestication significantly elevate in tumors compared to normal samples. Among the ten patients, the poor overlapping between the tumor-specific read-through or domestication events highlights the generally low frequency of these events, suggesting the deleterious effect conferred by these random processes. Since these tumor-specific read-through and domestication events are not observed in normal samples, and are likely to be avoided in healthy individuals, our study provides novel aspects in describing the landscape of oncogenesis, telling the readers that not only some particular genes are important for cancer, but also many particular mechanistic changes could be related to oncogenesis.

## Materials and Methods

### Data Availability

The transcriptome (RNA-seq) and translatome (ribosome footprints) data of ten patients of liver cancer were retrieved from the study (Zou et al., 2019). The accession number of data is GSE112705. The human and mouse reference genomes and the annotation of transposable elements were downloaded from UCSC genome browser. The genome versions are hg38 for human and mm10 for mouse. The cancer gene consortium (https://cancer.sanger.ac.uk/census/) contains the list of oncogenes and tumor suppressor genes. As said, the information of transposable element was contained in the reference genome website. The use of TE databases might be biased because the database could be over-complicated due to entangled sources (Bao et al., 2015), complex mechanisms and pipelines (Wei et al., 2016), or inaccessible websites (Roberg-Perez et al., 2003; Wu and Lu, 2019).

### Mapping the sequencing reads

We mapped the RNA-seq and ribosome profiling reads to the reference genome using hisat2 (Kim et al., 2019). The reads mapped to unique locations of the genome were kept. The tumor or normal samples from each individual were mapped separately. The IDs of the ten patients are LC001, LC033, LC034, LC501, LC502, LC505, LC506, LC507, LC508, and LC509 as provided by the original study. In our study, “genes with read-through events” means genes with at least one ribosome footprint mapped to the 3′UTR. Similarly, “domesticated TE” means the non-coding TE with at least one ribosome footprint detected. The locations of TE are given by the reference genome annotation. “Read-through ratio” for a gene is the percentage of ribosome footprints mapped to the 3′UTR to the footprints mapped to the whole gene region. For ribosome footprints in TE, a fraction is given for each TE as the number of ribosome footprints divided by the RNA reads mapped to this TE. The “tumor-specific read-through events” are defined as the genes with read-through events only appearing in tumor samples. Similarly, “tumor-specific protein domestication events” are the TEs that are specifically translated in tumor samples but not in normal samples.

### Random sampling of expressed genes

The RNA expression in each sample was provided by the original paper as RPKM values. To simulate the frequency spectrum of read-through or domestication events, we focus on the expressed genes (defined as RNA RPKM>0.1) in each sample. For example, assume that the numbers of genes expressed in ten patients are N1, N2, …, N10, and the numbers of genes with tumor-specific read-through events are n1, n2, …, n10. Of course, the n is smaller than N. The n1~n10 genes already have an observed frequency spectrum across the ten patients. Then, we randomly sample n1 genes from N1 genes, sample n2 genes from N2 genes, sample n10 genes from N10 genes. We look at the frequency spectrum of the randomly sampled genes, and compare this simulated result with the really observed frequency spectrum. The protein domestication (translated TE) is subjected to the same random sampling procedure.

### Mutations in mRNA-seq data

We used MuTect (do Valle et al., 2016) to treat the alignment files of mRNA-seq data. Default parameters were used but we ask the mutations to be derived mutations, that is, considering the ancestral state in mouse. The reference genome nucleotides should be identical for human and mouse for the mutation sites in the mRNA data. The matched positions and nucleotides were known from the alignments between human and mouse.

### Gene ontology enrichment

The GO analysis was conducted in R language with in-house packages.

### Graphic works

The graphic works were done in EXCEL or R platform.

## RESULTS

### Stop codon read-through is elevated in tumor samples

As we have introduced, the extent of stop codon read-through (potential protein extension) could be measured by the fraction of ribosome footprints in 3′UTRs. Theoretically, no ribosome footprints should be mapped to 3′UTRs. However, due to some inaccurate annotation of reference genome and the imperfect translation termination mechanism, there are always a small fraction of ribosome footprints mapped to the untranslated regions. We calculated the percentage of ribosome footprints in 3′UTRs (relative to the reads mapped to the whole gene region). In all the ten patients, this percentage is significantly higher in tumors than normal samples (Figure 2A). In normal samples, on average 3.2% of ribosome footprints were mapped to 3′UTRs, while in tumor samples, this average value is as high as 4.6%. In contrast, for RNA-seq data, roughly 48.2% of the reads could be mapped to 3′UTRs (relative to the whole gene region), with no difference between tumor and normal samples (Figure 2B). This is an indication that the higher ribosome density in 3′UTR of tumor samples is not caused by any potential biases in the mapping procedures.

**Figure 2.**
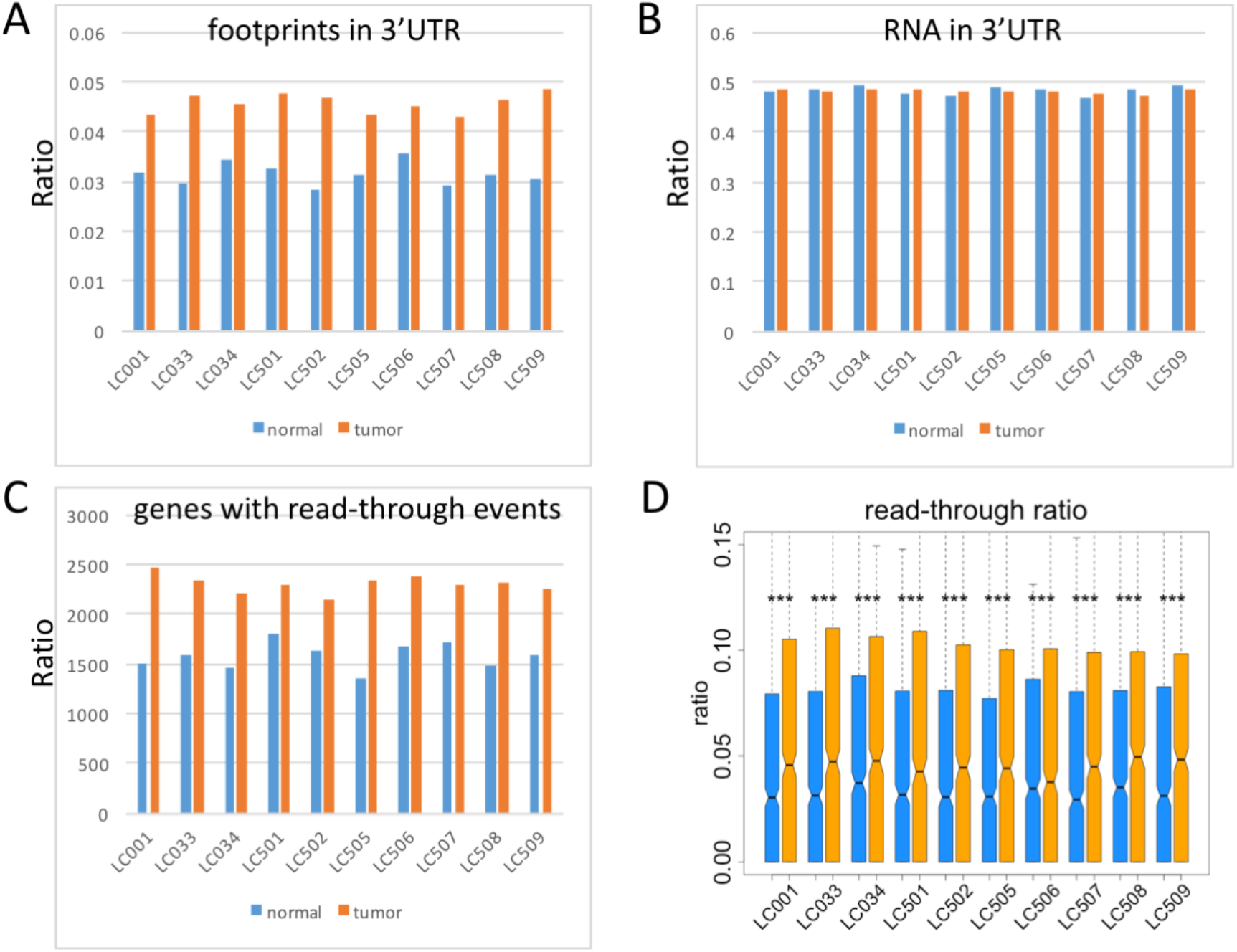
Protein extension (stop codon read-through) is elevated in tumor samples compared to normal samples. (A) The percentage of ribosome footprints mapped to the 3′UTR (relative to the whole gene region). The differences between tumor and normal samples are significant (proportion tests) for all ten individual patients (p-value < 1e-3 in all samples), and also significant for the “ten versus ten” non-parameter test (p-value = 0.0019). The IDs below the bars are the sample IDs used by the original study. (B) The percentage of RNA-seq mapped to the 3′UTR (relative to the whole gene region). None of the differences in the ten individuals are significant by using proportion tests. The “ten versus ten” non-parameter test is not significant either. (C) The numbers of genes with stop-codon read-through events. The differences between tumor and normal samples are significant (proportion tests) for all ten individual patients (p-value < 1e-3 in all samples), and also significant for the “ten versus ten” non-parameter test (p-value = 0.0019). (D) The read-through ratios per gene. “***” means p-value < 0.001 in the comparison between tumor and normal samples (KS tests).

After looking at the global ribosome footprints in 3′UTRs, which is an indicator of stop codon read-through, we are also curious about whether the global changes in read-through ratios are caused by the increased number of genes with read-through events or simply by the elevated read-through ratios of the same genes. We counted the numbers of genes with ribosome footprints detected in 3′UTRs (defined as genes with read-through events). We found that this number is obviously higher in all tumor samples compared to the matched normal samples (Figure 2C). On average, 1583 genes have detectable read-through events in normal samples while 2278 genes have detectable read-through events in tumor samples. Next, for each gene, we calculated the percentage of ribosome footprints in 3′UTRs (relative to the whole gene region) as their read-through ratios. This comparison also reveals globally higher read-through ratios of genes in tumors than normal samples (Figure 2D). We can conclude that the globally elevated read-through ratios in tumor samples are contributed by both the increased numbers of read-through genes and the increased power of read-through events in target genes.

### The tumor-specific read-through events are rare but enriched in oncogenes

We have shown that more genes bear stop codon read-through events in tumor samples than normal samples. We wonder whether these tumor-specific read-through events are deleterious. A reasonable prediction of deleterious events is that they should be observed with lower frequencies in population compared to neutral or beneficial events. We define the tumor-specific read-through events as the genes with read-through events only appearing in tumor samples. We calculate the frequency of the tumor-specific read-through events across ten patients. Meanwhile, we randomly sample the same numbers of genes with read-through events from the expressed gene pool in each individual. We find that the actual frequency spectrum is apparently skewed to low frequency compared to the simulated frequency spectrum (Figure 3A). To rule out potential technical biases, we calculate the occurrence of non-“tumor-specific” read-through events (shared between tumor and normal samples) and perform identical random sampling. The non-specific read-through events indeed show a slight tendency towards low frequency compared to simulated results (Figure 3B). However, the extent towards low frequency is much weaker for non-specific read-through events than the tumor-specific read-through events.

**Figure 3.**
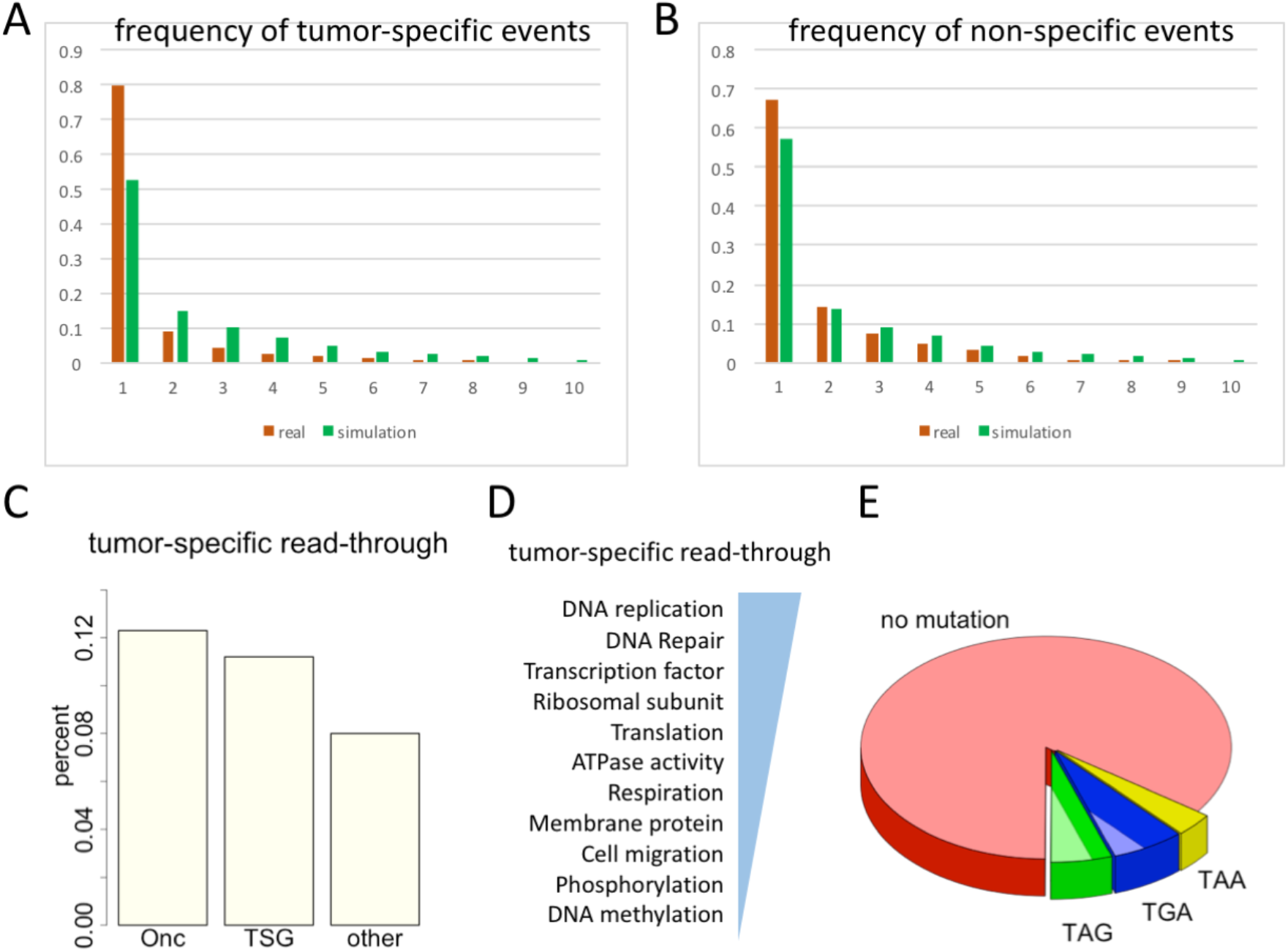
The frequency spectrum of read-through events across ten patients. (A) The observed (real) frequency spectrum and the simulated one. The tumor-specific read-through events are used. (B) The real frequency spectrum and the simulated one. The non-specific read-through events (those that are not unique to tumors) are used. (C) The percentages of read-through genes to be an oncogenes or tumor suppressor gene (TSG). (D) Gene ontology enrichment for the genes with tumor-specific read-through events. (E) For all the genes with tumor-specific read-through events, we check whether they have mutations on their stop codons. Only a small fraction of genes have mutated stop codons.

Notably, although the tumor-specific read-through events themselves are mostly singletons across the ten patients, these events have higher occurrence in oncogenes and tumor suppressor genes (TSG) compared to the remaining genes (Figure 3C). Roughly less than 10% of expressed genes have tumor-specific read-through events, but among oncogenes and TSG, this fraction is higher than 10%. It is known that oncogenes play positive roles in oncogenesis while TSG play counteractive roles. However, the read-through event in a gene does not necessarily enhance/promote or reduce/damage its function, the read-through only potentially extends the protein sequence. This protein extension could be either deleterious or beneficial (for a gene), but for the individual patient, the effect might be harmful since these tumor-specific read-through events are present in tumor and absent in normal samples.

For the genes with tumor-specific read-through events, we exclude the oncogenes and TSG and perform gene ontology enrichment analysis. It turns out that these genes are enriched in various important cellular pathways like DNA replication, mRNA transcription and translation (Figure 3D). This gives a reason that the extended protein caused by read-through events might have a great impact on the entire cellular processes, making these abnormal translation events deleterious. Next, we try to look for potential factors responsible for the seemingly random read-through events. We can attribute a small fraction of read-through events to the mutations on stop codons revealed by RNA-seq data, however, for the majority of genes with read-through events, no mutations are observed on the stop codons (Figure 3E). This again suggests the randomness of the abnormal read-through events, which could be caused by multiple undetermined factors.

### The protein domestication from non-coding region is prevalent in tumor samples

As we have introduced, both protein extension by read-through events and protein domestication from non-coding repetitive elements could diversify the cellular protein pool. The changes caused by the two mechanisms could either be beneficial or deleterious, which depends on the function of target genes. Here we perform very similar analyses as we did in previous parts. In summary, we find that the fractions of ribosome footprints mapped to the transposable elements (TEs that are non-coding, footprints relative to RNA) are higher in tumor samples than normal samples, globally (Figure 4A) or individually (Figure 4B). The numbers of transposable elements with footprints detected (termed protein domestication events) are higher in tumor samples than normal samples (Figure 4C). Finally, the frequency spectrum across ten patients suggests that the tumor-specific protein domestication events (the TEs specifically translated in tumors but not normal samples) are skewed to low frequency (Figure 4D) and seem to be deleterious for the individuals.

**Figure 4.**
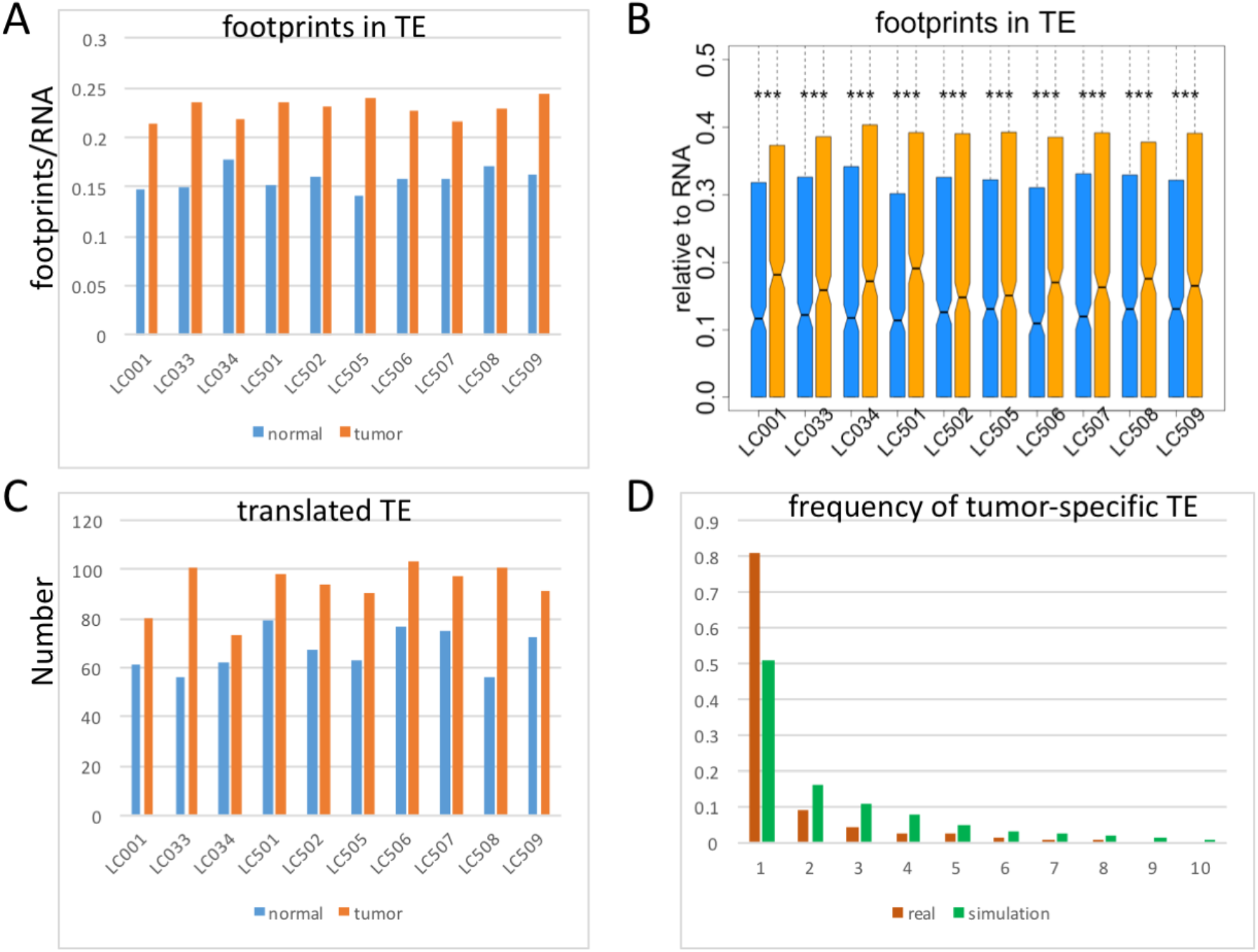
The protein domestication (translated TE) events. (A) Fraction of ribosome footprints mapped to non-coding TE regions normalized by RNA reads. The difference is significant for the “ten versus ten” non-parameter test (p-value = 0.0019), and also significant for individually testing the two fractions in each sample (p-value < 1e-3 in all samples). (B) Fraction of ribosome footprints mapped to each TE normalized by RNA reads. “***” means p-value < 0.001 by using KS tests. (C) Numbers of translated TEs in normal and tumor samples of each individual patient. The difference is significant for the “ten versus ten” non-parameter test (p-value = 0.0019), and also significant for individually testing the two fractions in each sample (p-value < 1e-3 in all samples). (D) Frequency spectrum of the tumor-specific translated TE. The real and simulated frequencies are shown.

Since these patterns are highly identical to those observed for stop codon read-through (protein extension) events, we can conclude that the tumor-specific protein extension and domestication events correlate with liver cancer oncogenesis, and are likely to play a role during the cancer development process. These events are of low frequency among the ten patients, and may otherwise be purged from the population if not sequenced timely.

### Mutation profile reveals possible role of the protein extension and domestication in oncogenesis

We are curious whether we can dig out any clues from the mutation profile of these samples. The mutations in a sample could be known from the mRNA-seq data. Although the traditional mutation calling pipeline requires genome re-sequencing, here we only care about the mutations in coding regions and UTRs, and therefore the mRNA data are deep enough to accomplish this. We ask a mutation in the patient sample to be derived mutation by checking the matched genomic position in mouse. Only the mutation sites with identical nucleotide in human and mouse reference genomes are kept.

One basic assumption is, if the read-through or TE domestication events really contribute to oncogenesis, then the tumor-specific events should be advantageous and the mutations in those regions should be positively selected in tumor sample (although they are deleterious to the individual patients). The selection should be reflected by the enrichment of mutations in the “effective regions” of these events, which are, 3′UTRs for read-through events and coding regions for TE domestication events.

We divide all read-through genes into tumor-specific and non-specific genes. We count the numbers of mutations in 3′UTRs and other regions. The fraction of mutations in 3′UTR is higher in tumor-specific genes and lower in non-specific genes (Figure 5A). We divide all domesticated TEs (with ribosome footprints detected) into tumor-specific and non-specific TEs. The fraction of mutations in CDS is higher in tumor-specific TEs and lower in non-specific TEs (Figure 5B). These results verify our assumption that the tumor-specific events might play a role in oncogenesis as the mutations are enriched in the “effective regions” of these events (3′UTRs for read-through events and coding regions for TE domestication events).

**Figure 5.**
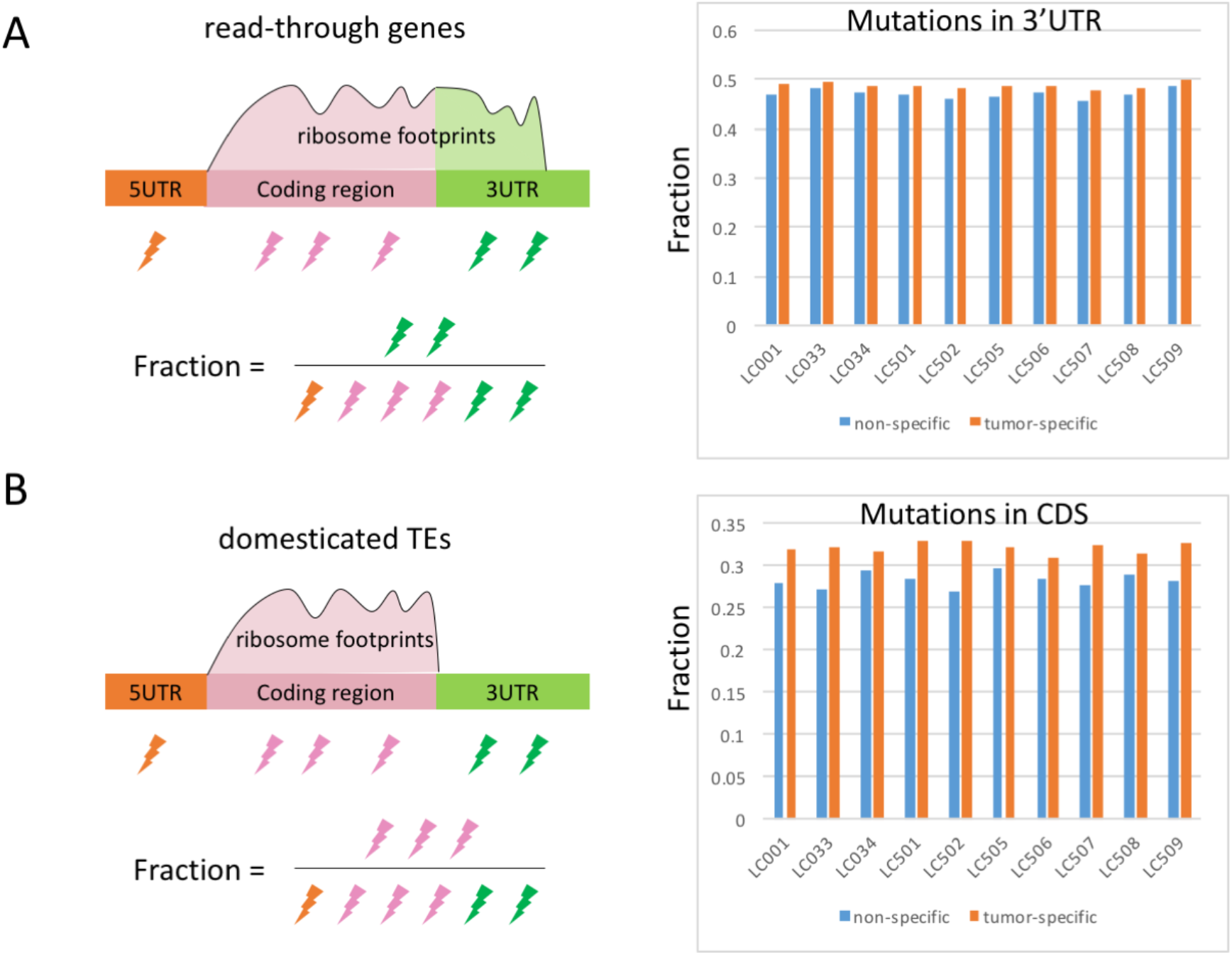
Mutation profiles of read-through genes and domesticated TEs. (A) The read-through genes are divided into tumor-specific genes and non-specific genes. We calculate the fraction of mutations in 3′UTR regions. The difference is significant for the “ten versus ten” non-parameter test (p-value = 0.0019), but insignificant for individually testing the two fractions in each sample (p-value > 0.05 in all samples). (B) The domesticated TEs are divided into tumor-specific genes and non-specific genes. We calculate the fraction of mutations in coding regions. The difference is significant for the “ten versus ten” non-parameter test (p-value = 0.0019), and also significant for individually testing the two fractions in each sample (p-value < 1e-2 in all samples).

## DISCUSSION

While the gene by gene examination could be accomplished by experimental biologists, the mechanism by mechanism examination is the gift from the big data era. The advantage of high-throughput sequencing data is that it allows us to look into any cis-regulatory changes at genome-wide level. For example, when testing whether the read-through ratios are different between tumor and normal samples, the rejection of null hypothesis would indicate the global change in a mechanism rather than the change in particular genes. Although some mutations in particular genes might account for the observed read-through events, the overall tendency is to show the mechanistic change. There is high confidence to claim that the occurrence of stop codon read-through events is elevated in tumor sample, however, for any specific genes, it is hard to give a probability of whether this gene has significantly changed its read-through ratio. To test for specific genes, it would better to design cellular experiments to mimic the in vivo situation.

Natural selection favors particular changes and elevates their frequencies in a population. For the deleterious mutations, natural selection suppresses them to very low frequencies or directly sweeps them out, termed extinction. Therefore, extinction events are taking place all the time. The tumor samples are precious materials to study evolution because the patients carrying the deleterious changes (or mutations) are almost on the edge of extinction (no offense). Once eliminated, the changes or mutations would never have the chance to be observed. The tumor samples allow us to trace back in time and look for the patterns of deleterious changes.

Now that we have observed globally higher protein extension and domestication events in tumor samples than normal samples, suggesting that these events might correlate with oncogenesis. Ironically, although the tumor-specific protein extension and domestication events might facilitate the formation of tumor (they could be beneficial for the tumor cells), they are definitely deleterious to the individual patients. The frequency of occurrence across the ten patients further supports the harmful effect of these events.

One more issue to be discussed is the potential causal factors of protein domestication events. In the above sections of the protein extension events, we have already tried to explain (a small part of) the stop codon read-through events by the mutations that abolish stop codons. The majority of the read-through events are still to be explained by randomness, though. Here, for the protein domestication (TE domestication) events, we will discuss why some of the TEs are translated (with ribosome footprints) and some are not. We assume that different coding sequences have different coding potential. TEs with high coding potential CDS should be preferentially translated, becoming a domesticated TE. Nest, we suppose that the actively transcribed TEs are easier to encounter free ribosomes and therefore have higher probability to be translated and domesticated. Besides, the length of a TE (and the length of its protein) may also decide how worthy it is to take a risk (energy waste during translating) to try on this new TE. All these factors plus randomness could affect the choice of which TEs to be translated and domesticated. Not testing these factors is also the limitation of our work. To fix this limitation in future studies, we need to analyze and compare these features in domesticated and non-domesticated TEs. A clear difference should be seen if the proposed factors really contribute to TE domestication.

Our study also re-emphasizes the notion that one gene does not represent one sequence. One gene could be transcribed to different RNA isoforms, and then produce various forms of protein sequences. Unfortunately, this notion did not receive sufficient attention. In many cancer studies, the differential expression analyses of single cell sequencing data are over-simplified because they do not consider the isoform changes or any other “intra-gene” changes (like stop codon read-through which replenishes the cellular pool). Indeed, the single cell sequencing might not be deep enough for a cell to analyze the isoform changes. However, the technical limitation is not the reason to ignore this concern. Particular examples could show that there might be false discoveries of the differentially expressed genes. Assume that gene B expresses B1 isoform in normal samples and B2 isoform in tumor samples. In both conditions, there are 100 molecules. If B2 is longer than B1, then there should be more sequencing reads in tumor (B2 isoform) than in normal samples (B1 isoform). Gene B would be erroneously regarded as differentially expressed genes. But actually, both conditions have 100 molecules (in other words, they are not differentially expressed at all), the only difference is that isoform B2 is longer than isoform B1. This fact promotes the fact that the precise medicine claimed by the single cell sequencing biotechnology is actually not precise. There are still spaces for improvement like increasing the sequencing depth to allow deeper analyses on isoform changes and other “intra-gene” changes.

In conclusion, the advantages of our work are, (1) We find the interesting patterns of elevated protein extension (stop codon read-through) and domestication (translating TE) events in tumor samples compared to normal samples. Both types of events diversify the cellular pool; (2) We surmise that the tumor-specific read-through or domestication events are deleterious to the individual patients, with support from the observed frequency spectrum; (3) We propose the idea that not only particular genes should be examined, but also the particular mechanisms should be investigated in the cancer studies; (4) One gene does not refer to only one sequence. Many changes at the RNA or protein levels could increase the sequence pool of the same gene. Therefore, merely looking for the differentially expressed genes between tumor and normal tissues is not enough to reflect the real landscape of oncogenesis. This importance issue should be paid adequate attention to and should no longer be ignored by this field.

## Abbreviations

mRNA: messenger RNA.
TE: transposable elements.
TSG: tumor suppressor gene.
UTR: untranslated region.
CDS: coding sequence or coding region.
RPKM: reads per kilobase per million mapped reads.

## Acknowledgements

We thank our colleague Dr. Wang for the guidance on graphical works. We thank our institution for the support during this COVID-19 time. In memory of Xiaolei Wang (Wang et al., 2018), as we all know, and Liming Wang for the comments on this.

## Author contributions

All authors approved this manuscript. The corresponding author supervised this whole project.

## Funding

No funding has supported this research.

## Compliance with ethical standards

### Conflict of interest

The authors declare that they have no conflict of interest.

### Ethical approval

This article does not contain any studies with human participants or animals performed by any of the authors.

### Data availability

The transcriptome and translatome data of ten patients of liver cancer were retrieved from the study (Zou et al., 2019). The accession number of data is GSE112705. The human reference genome was downloaded from UCSC genome browser under version hg38.

## Notes

### Competing Interest Statement

The authors have declared no competing interest.

## References

Bagger, F.O., and Probst, V. (2020). Single Cell Sequencing in Cancer Diagnostics. Adv Exp Med Biol 1255, 175–193.

Bao, W., Kojima, K.K., and Kohany, O. (2015). Repbase Update, a database of repetitive elements in eukaryotic genomes. Mob DNA 6, 11.

Chandola, U., Das, R., and Panda, B. (2015). Role of the N6-methyladenosine RNA mark in gene regulation and its implications on development and disease. Brief Funct Genomics 14, 169–179.

do Valle, I.F., Giampieri, E., Simonetti, G., Padella, A., Manfrini, M., Ferrari, A., Papayannidis, C., Zironi, I., Garonzi, M., Bernardi, S. et al. (2016). Optimized pipeline of MuTect and GATK tools to improve the detection of somatic single nucleotide polymorphisms in whole-exome sequencing data. BMC Bioinformatics 17, 341.

Fung, T.S., and Liu, D.X. (2018). Post-translational modifications of coronavirus proteins: roles and function. Future Virol 13, 405–430.

Kim, D., Paggi, J.M., Park, C., Bennett, C., and Salzberg, S.L. (2019). Graph-based genome alignment and genotyping with HISAT2 and HISAT-genotype. Nat Biotechnol 37, 907–915.

Kochetov, A.V., Sarai, A., Rogozin, I.B., Shumny, V.K., and Kolchanov, N.A. (2005). The role of alternative translation start sites in the generation of human protein diversity. Mol Genet Genomics 273, 491–496.

Liu, J., Chang, H.W., Huang, Z.M., Nakamura, M., Sekhon, S., Ahn, R., Munoz-Sandoval, P., Bhattarai, S., Beck, K.M., Sanchez, I.M., et al. (2020). Single cell RNA-seq of psoriatic skin identifies pathogenic Tc17 subsets and reveals distinctions between CD8(+) T cells in autoimmunity and cancer. J Allergy Clin Immunol.

Liu, Q., Hu, H., and Wang, H. (2015). Mutational bias is the driving force for shaping the synonymous codon usage pattern of alternatively spliced genes in rice (Oryza sativa L.). Mol Genet Genomics 290, 649–660.

Magistri, M., Velmeshev, D., Makhmutova, M., and Faghihi, M.A. (2015). Transcriptomics Profiling of Alzheimer’s Disease Reveal Neurovascular Defects, Altered Amyloid-beta Homeostasis, and Deregulated Expression of Long Noncoding RNAs. J Alzheimers Dis 48, 647–665.

Murray, J.S., and Murray, E.H. (2016). TE-domestication and horizontal transfer in a putative Nef-AP1mu mimic of HLA-A cytoplasmic domain re-trafficking. Mob Genet Elements 6, e1176634.

Okada, S., and Brennicke, A. (2006). Transcript levels in plant mitochondria show a tight homeostasis during day and night. Mol Genet Genomics 276, 71–78.

Roberg-Perez, K., Carlson, C.M., and Largaespada, D.A. (2003). MTID: a database of Sleeping Beauty transposon insertions in mice. Nucleic Acids Res 31, 78–81.

Sbodio, J.I., Snyder, S.H., and Paul, B.D. (2016). Transcriptional control of amino acid homeostasis is disrupted in Huntington’s disease. Proc Natl Acad Sci U S A 113, 8843–8848.

Soong, B.W., Hsieh, S.Y., and Chak, K.F. (1994). Mapping of transcriptional start sites of the cea and cei genes of the ColE7 operon. Mol Gen Genet 243, 477–481.

Tieng, F.Y.F., Baharudin, R., Abu, N., Mohd Yunos, R.I., Lee, L.H., and Ab Mutalib, N.S. (2020). Single Cell Transcriptome in Colorectal Cancer-Current Updates on Its Application in Metastasis, Chemoresistance and the Roles of Circulating Tumor Cells. Front Pharmacol 11, 135.

Wang, X., Chen, Z.H., Yang, C., Zhang, X., Jin, G., Chen, G., Wang, Y., Holford, P., Nevo, E., Zhang, G., et al. (2018). Genomic adaptation to drought in wild barley is driven by edaphic natural selection at the Tabigha Evolution Slope. Proc Natl Acad Sci U S A 115, 5223–5228.

Wei, G., Qin, S., Li, W., Chen, L., and Ma, F. (2016). MDTE DB: a database for microRNAs derived from Transposable element. IEEE/ACM Trans Comput Biol Bioinform 13, 1155–1160.

Wu, C., and Lu, J. (2019). Diversification of Transposable Elements in Arthropods and Its Impact on Genome Evolution. Genes (Basel) 10.

Zou, Q., Xiao, Z., Huang, R., Wang, X., Wang, X., Zhao, H., and Yang, X. (2019). Survey of the translation shifts in hepatocellular carcinoma with ribosome profiling. Theranostics 9, 4141–4155.

